# Wood modification by furfuryl alcohol caused delayed decomposition response in *Rhodonia (Postia) placenta*

**DOI:** 10.1101/454868

**Authors:** Inger Skrede, Monica Hongrø Solbakken, Jaqueline Hess, Carl Gunnar Fossdal, Olav Hegnar, Gry Alfredsen

## Abstract

The aim of this study was to investigate differential expression profiles of the brown rot fungus *Rhodonia placenta* (previously *Postia placenta*) harvested at several time points when grown on *Pinus radiata* (radiata pine) and *P. radiata* with three different levels of modification by furfuryl alcohol, an environmentally benign commercial wood protection system. For the first time the entire gene expression pattern of a decay fungus is followed in untreated and modified wood from initial to advanced stages of decay. Results support the current model of a two-step decay mechanism, with an initial oxidative depolymerization followed by hydrolysis of cell-wall polysaccharides. The wood decay process is finished, and the fungus goes into starvation mode after five weeks when grown on unmodified *P. radiata* wood. The pattern of repression of oxidative processes and oxalate synthesis found in *P. radiata* at later stages of decay is not mirrored for the high furfurylation treatment. The high treatment level provided a more unpredictable expression pattern throughout the entire incubation period. Furfurylation does not seem to directly influence the expression of core plant cell wall hydrolyzing enzymes, as a delayed and prolonged, but similar pattern was observed in the *P. radiata* and the modified experiments. This indicates that the fungus starts a common decay process in the modified wood, but proceeds at a slower pace as access to the plant cell wall polysaccharides is restricted. This is further supported by the downregulation of hydrolytic enzymes for the high treatment level at the last harvest point (mass loss 14%). Moreover, the mass loss does not increase the last weeks. Collectively, this indicates a potential threshold for lower mass loss for highly modified wood.

**IMPORTANCE:** Fungi are important decomposers of woody biomass in natural habitats. Investigation of the mechanisms employed by decay fungi in their attempt to degrade wood is important for both the basic scientific understanding of ecology and carbon cycling in nature, and for applied uses of woody materials. For wooden building materials long service life and carbon storage is essential, but decay fungi are responsible for massive losses of wood in service. Thus, optimizing durable wood products for the future are of major importance. In this study we have investigated the fungal genetic response to furfurylated wood, a commercial environmentally benign wood modification approach, that improves service life of wood in outdoor applications. Our results show that there is a delayed wood decay by the fungus as a response to furfurylated wood and new knowledge about the mechanisms behind the delay is provided.

## INTRODUCTION

Wood as a building material has a number of attractive properties including carbon sequestration during its service life and its aesthetic aspects as a natural material. One of the main challenges with using wood as a building material is its susceptibility to attack by wood degrading microorganisms. Traditional wood preservatives with a biocidal mode of action (e.g. organic- or copper-based preservatives) cause concerns due to their perceived negative environmental impacts. In Europe the Construction Products Regulation and the Biocidal Products Regulation have a major impact on the wood industry and the number of active ingredients allowed for wood preservatives is decreasing (1). This, together with new consumer awareness and demand for more environmentally focused products (2), pushes the need for new products and wood protection approaches.

The modern wood modification approach, in contrast to more traditional wood preservatives, is to modify the wood matrix so that it can no longer act as a suitable substrate for wood degrading organisms. Different wood modification approaches were described in detail by Hill (3). Currently the processes underlying available wood modification products can be classified as chemical processing (acetylation, furfurylation, resin impregnation etc.), thermo-hydro processing (thermal treatment) and thermo-hydro-mechanical processing (surface densification) (2). The different wood modification processes are at various stages of development, and the most established processes on the marked include thermal treatment, acetylation and furfurylation.

The commercial wood modification process furfurylation, use furfuryl alcohol which is manufactured industrially by catalytic reduction of furfural obtained from agricultural waste such as sugar cane bagasse or corn cobs (4). It involves a wood impregnation step with furfuryl alcohol and catalysts, followed by a curing step where the furfuryl alcohol is polymerized within the wood cell walls (2, 4). This is a complex chemical reaction and there is still some discussion whether the furfurylation process only bulks the wood cell wall or if it also causes chemical modifications of the native wood cell wall polymers. As support for the latter theory it has been shown in a model lignin system that the furfural polymer formed covalent bonds with lignin (5). In addition to this uncertainty, the mechanisms utilized by decay fungi in their attempt to degrade modified wood are not well understood at the molecular level and this hinders knowledge-based design of the modification methods.

Independent of how the furfurylation modifies the wood, the process results in improved resistance to fungal deterioration (2-4, 6, 7). The level of modification by furfural alcohol is measured by the Weight Percent Gain (WPG) of the wood. Lande et al. (6) concludes that furfurylated wood treated to a WPG of 35% or more, has sufficient resistance to brown and white rot decay fungi.

Traditionally wood degrading basidiomycetes have been divided into white- or brown rot fungi due to the ability of white rot fungi only to degrade lignin along with holocellulose, while brown rot fungi leave the lignin behind as a brown residue. White rot fungi have a larger repertoire of known enzymes that depolymerize the components of the plant cell wall than brown rot fungi (8). It is important to note that this traditional dichotomy is polyphyletic and it has been suggested, based on comparative studies of 33 basidiomycetes, that a continuum exists, without a clear distinction between the types of decay (8). Brown rot fungi dominate decomposition of conifer wood in boreal forests even if only 6% of fungal wood decay species produce the classical brown rot decay (9). Moreover, brown rot causes more challenges than white rot for wood in service outdoors (10). Brown rot fungi depolymerize cellulose rapidly during incipient stages of wood colonization, resulting in considerable losses in strength even at early decay stages (11, 12). The current hypothesis is that brown rot fungi utilize a non-enzymatic system that rapidly depolymerizes cell wall components in early stages of decay, prior to depolymerization by traditional cellulases and hemicellulases (13-15). This system is often referred to as the Chelator Mediated Fenton (CMF) system and involves the solubilization of iron from the environment by oxalic acid that the fungus secretes, followed by the reduction of iron by chelating/reducing secondary metabolites that then can react with hydrogen peroxide within the wood cell wall. It is theorized that this produces oxygen radicals through a Fenton-like reaction that will depolymerize lignocellulose and make these polymers available for large hydrolytic enzymes (16). This two-step theory has been further supported by the gene expression pattern in *Rhodonia placenta* (17). Whether the CMF functions as a pretreatment of the wood opening the cell walls to allow the enzymes to enter the cell wall, or whether polysaccharide components diffuse into the cell lumen to the enzymes is currently debated (18). However, there is agreement in that the brown rot CMF systems and the hydrolytic enzymes cannot be present at the same time since highly reactive ROS (reactive oxygen species) reactions will cause oxidative damage on glycoside hydrolases. Zhang and Schilling (19) found that cellobiose appears to play a key role in the transition between the oxidative phase and the hydrolytic phase. However, the full extent of feedback mechanisms regulating brown rot decay is not known.

*Rhodonia* (*Postia*) *placenta* is a commonly used brown rot decay fungus in laboratory wood decay tests, strain FPRL 280 is included the European standard EN 113 (20) and strain ATCC11538 is included in the American standard E10-16 (21). The *R. placenta* strain MAD 698-R, and its monokaryotic strain MAD698-R-SB12 has been genome sequenced by the Department of Energy, Joint Genome Institute (JGI), USA (22, 23). The species has frequently been used as a model fungus for gene expression studies of untreated wood (17, 19, 23-25) and modified wood (26-32), including furfurylated wood (28, 31). These experiments have shown that several hemicellulases but few potential cellulases were produced when the fungus was grown on ball-milled aspen or glucose as substrate (24). The expression patterns for oxidoreductase encoding genes support an extracellular Fenton system. Furthermore, Skyba et al. (33) demonstrated that the gene expression profile of *R. placenta* (and *Phanerochaete chrysosporium*) was influenced by wood substrate composition (three *Populus trichocarpa* genotypes) and the duration of incubation. For early stages of modified wood, a possible shift toward increased expression of genes related to oxidative metabolism and concomitant reduction of several gene products related to the breakdown of holocellulose in furfurylated wood compared to unmodified wood has been suggested (28).

In this study we investigated the differential expression profiles of the brown rot fungus *Rhodonia placenta* harvested at several time points when grown on *Pinus radiata* with three different levels of furfurylation. For comparison we also investigated the gene expression during decay of unmodified *P. radiata*. The overall aim is to understand the mechanisms utilized by brown rot decay fungi in their attempt to degrade modified and unmodified wood. This is important for further optimization of future modified wood products, and for an expanded understanding of the fungal decay process in general.

## RESULTS

In the following experiments the *P. radiata* wood was modified at three different levels. The three modification levels had a mean Weight Percent Gain (WPG) of 3.8±0.7%, 24.0±3.5%, 36.6±5.0%. For simplicity the treatments are named WPG4, WPG24 and WPG37, respectively. The experiment on the unmodified wood is named *P. radiata*.

### Mass loss calculations

Five weeks after inoculation of *R. placenta* strain FPRL 280 on the *P. radiata* wood blocks, they had a mean mass + treatment loss of 28.8±4.0%, while the modified WPG37 only had a mean mass + treatment loss of 13.5±3.3% after 21 weeks (Fig. 1). Mass loss is within the wood protection literature mostly referred to as the mass loss of the entire wood + treatment system, i.e. dry weight of the treated wood before fungal inoculation compared to dry weight after fungal decay. Another way to measure mass loss is to assume that only the wood is decayed and eliminate the WPG of the treatment from the mass loss calculation. Both methods resulted in the same general trends, but the exclusion of WPG results in slightly higher mass losses as expected (Fig. S1). The differences between the two mass loss calculation approaches for the three furfurylation levels at the last harvesting points were (wood + treatments vs. only wood): WPG4 week 9 41.7% vs. 42.5%, WPG24 week 21 29.0% vs. 34.6%, WPG37 week 21 13.5% vs. 16.0%. We use the wood + treatment results as presented in Fig. 1 as our mass loss detection system.

**Figure 1.**
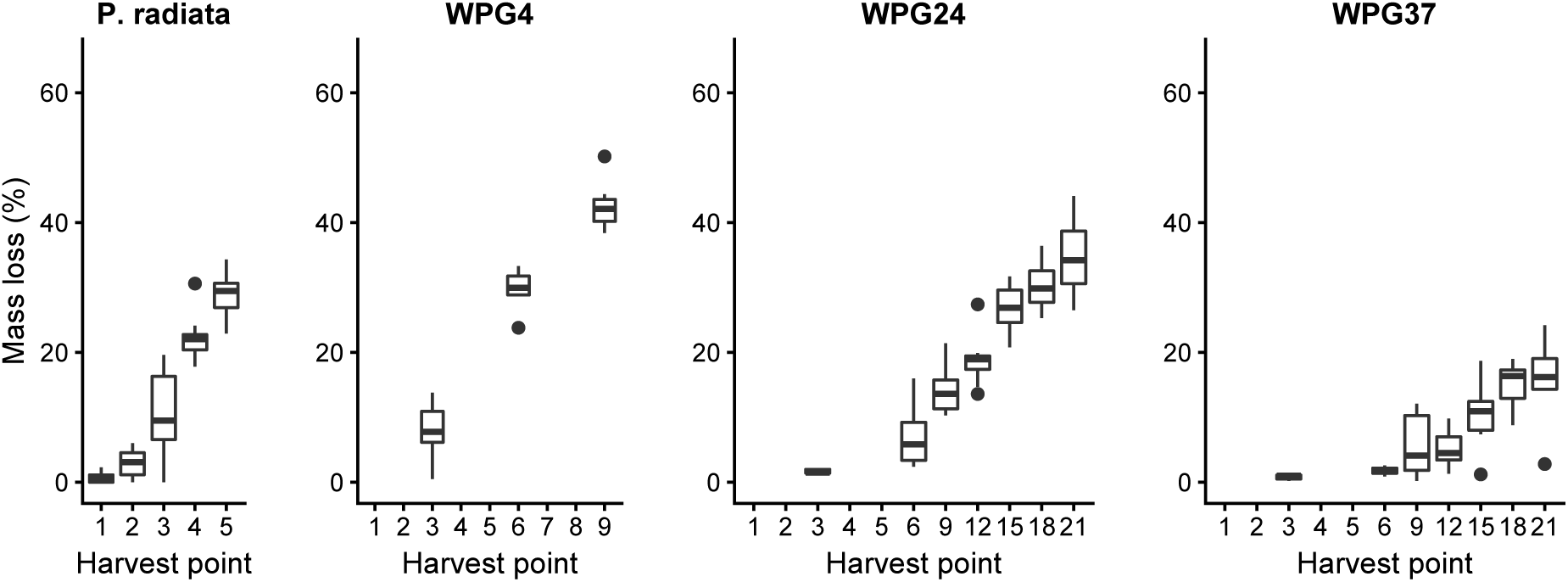
Boxplots of the mass loss of all experiments of *Rhodonia placenta* grown on *Pinus radiata* and different levels of modification by furfurylated *P. radiata*. The weight of the treatment is included the measurements. a) *R. placenta* grown on unmodified P. radiata. Wood harvested at five different harvest points (week). b) *R. placenta* grown on furfurylated *P. radiata*, Weight Percent Gain (WPG) 4%. Wood harvested at three different harvest points. For c) and d) The wood was harvested at six harvest points for *R. placenta* grown on furfurylated *P. radiata*, WPG 23% (c) and WPG 37% (d).

### Differential gene expression

In order to investigate the genetic basis of the different behavior of *R*. *placenta* growing on different levels of furfurylated wood we sequenced transcriptomes of a selection of time series using Illumina Nextseq sequencing. The sequenced time points for each treatment were sampled according to the following setup: *P. radiata* – weeks 1, 2, 3, 4 and 5, WPG4 – weeks 3, 6 and 9, WPG24 – weeks 3, 6, 9, 12, 15, and 18, WPG37 – weeks 3, 6, 9, 12, 15 and 18.

Even if the genome and transcriptome of the American *R. placenta* strain MAD 698-R has been sequenced, this could not be used for mapping purposes in this study. The genome sequenced American strain MAD 698-R and the European strain FPRL 280 used in this study are significantly different, with a mapping success of only 40-60% when the reads were mapped to the genome of MAD 698-R (results not shown). We therefore produced Illumina Hiseq paired-end data to be used for a transcriptome assembly. The resulting assembly had 56 520 contigs (transcripts), and covered 99.3% of the conserved BUSCO fungal genes (Table S1; Table S2). In the transcriptome, 9 355 contigs had an annotation from Blastx, and 18 917 contigs were given a PFAM annotation. In all further analyses the sequence data were mapped to this annotated transcriptome assembly. The mapping success of sequence data to this transcriptome was more than 90% for all libraries.

The gene expression data demonstrate consistent results for each treatment with a gradient clustering of replicate samples according to treatment with few outliers (Fig. 2). There is some biological variation between replicates as expected from natural variation in wood and the variation in modification of the wood blocks. The *P. radiata* and WPG37 experiments display a wider range of biological variation compared WPG24 and WPG4.

**Figure 2.**
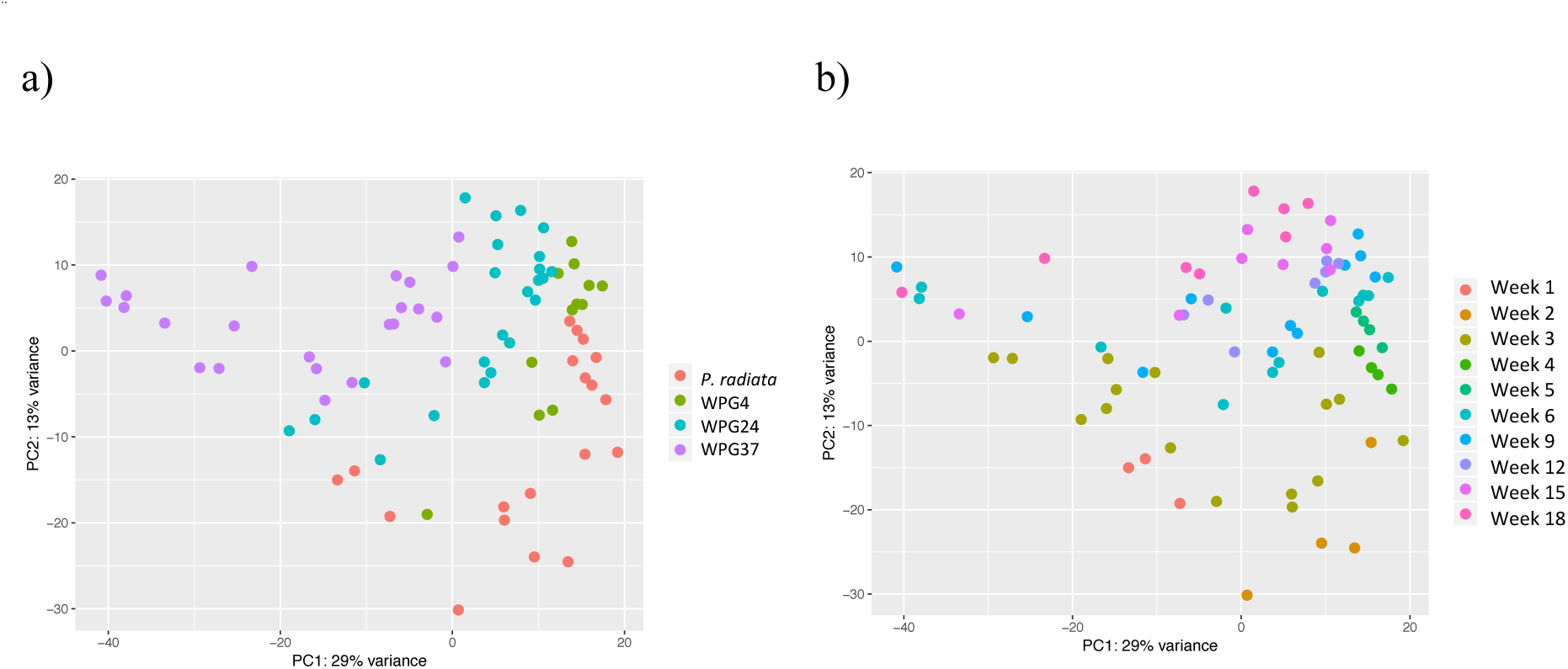
The figures of PCA plots of all the RNAseq replicates of *Rhodonia placenta* grown on *Pinus radiata* and furfurylated *P. radiata*. In total 42% of the variance is explained by the two PCA axis. a) and b) show the same plot, but in a) the replicates are colored according to experiment and in b) the replicates are colored according to harvest time (week).

### Differential gene expression and Functional enrichment across treatments

To obtain an overall impression of the entire dataset we ran a multifactorial DE analysis, factoring in time and treatment. Thus, all harvest time points were included for all treatments and we extracted contrasts describing the different modified treatments versus the unmodified *P. radiata* experiment, i.e. WPG4 vs *P. radiata*, WPG24 vs *P. radiata* and WPG37 vs *P. radiata*. (Table 1). Functional enrichment analyses of the resulting gene lists were inferred using the annotated PFAM domains and GO terms from the annotated transcriptome. Compared to *P. radiata*, the WPG4 showed upregulation of zinc-binding dehydrogenase domains (PF00107.21) and two GO terms related to zinc ion binding and oxidoreductase activity. The same enrichment was also found upregulated in WPG24 and WPG37 when compared to *P. radiata*. In addition, these two latter treatments showed upregulation of more oxidoreductase domains, most in WPG37. No PFAM domains or GO terms were found downregulated in WPG4 or WPG24. However, WPG37 showed a strong downregulation of eight PFAM domains and seven GO terms with functions related to protein and peptide degradation, the ubiquitin-proteasome pathway and the Ras gene family compared to *P. radiata* (Table 1), e.g. two domains related to proteasome (PF10584.4 and PF00227.21) and Ras (PF08477.8 and PF00071.17). These terms were not found among the other comparisons.

**Table 1.**
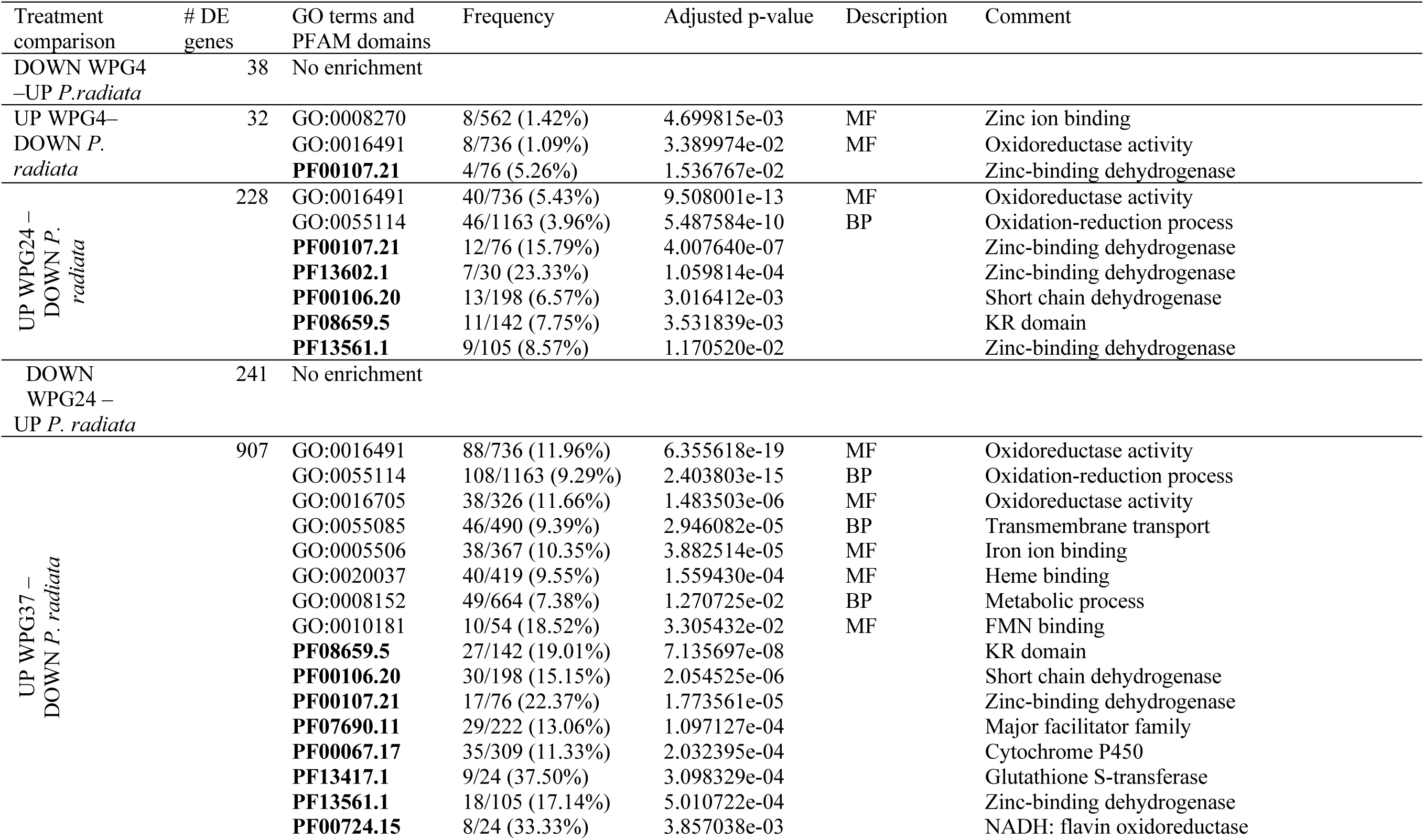
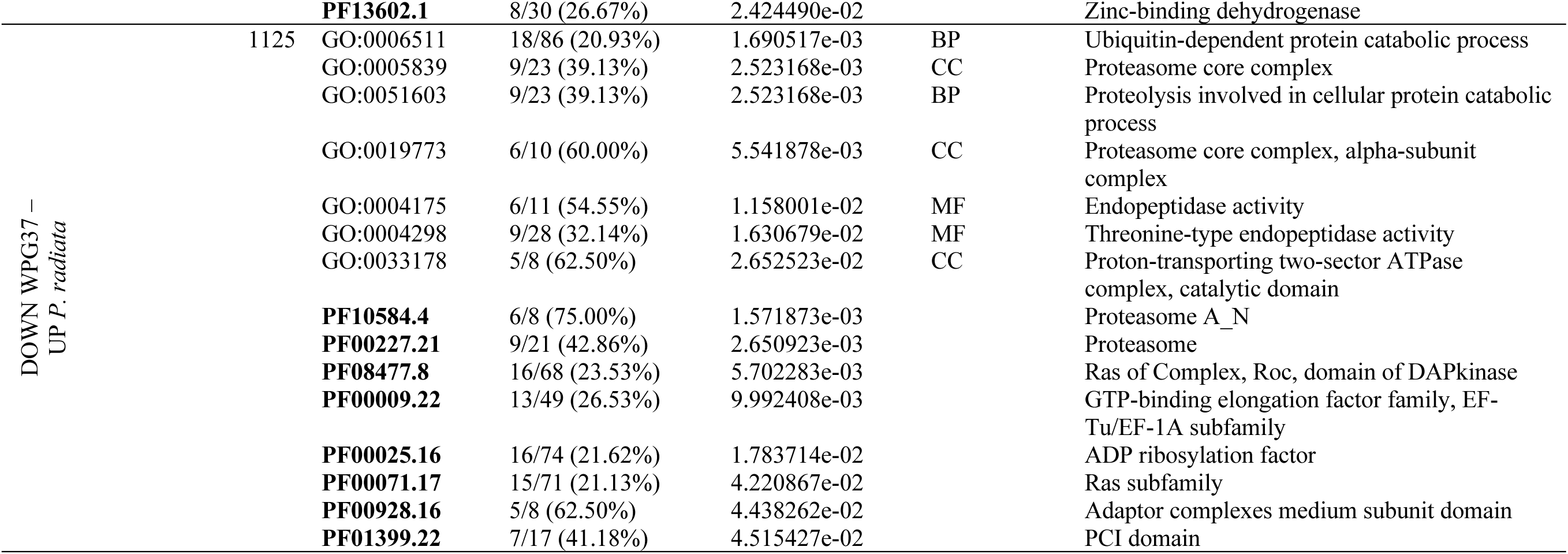
Functional enrichment analyses of GO terms and PFAM domains of significant differential expressed genes between treatments along the time series of *Rhodinia placenta* growing on unmodified and three different levels of modification with furfuryl alcohol (Weight Percent Gain (WPG) of wood of 4, 24 and 37%). Description indicates gene ontology variables; MF – Molecular function, BP – Biological process.

As an alternative analysis method to the multifactorial DE analyses above, we also clustered the genes with similar expression profiles. The read counts were grouped into ten clusters (K) for each treatment, and samples from *P. radiata*, WPG4, WPG24 and WPG37 were analyzed separately (Fig. 3). For these clusters we did functional enrichment with PFAM and GO terms (Table S3), and also investigated the placement of known genes related to plant cell wall decay in these clusters (Table S4).

For most of the clusters, an observed pattern of higher expression in one or a few of the harvest points was found (Fig. 3). For *P. radiata*, two clusters were directly linked to wood decay and carbohydrate active enzymes (Table S3). One of these clusters (*P. radiata*-K2) was related to early depolymerization of hemicellulose and pectin (enriched for GH28 and GH43 domains and containing the genes OxaD, Man5a and CE16b; see Table 2 for information about abbreviated gene names) and highly expressed in week 1, while the other cluster (*P. radiata*-K5) was related to later stages of cellulose depolymerization (enriched for GH3 and hydrolase activity, and containing the genes Cel5b, Cel2, bGlu, Xyl10a, bXyl; Table 2) and was highly expressed in week 2. Similar enrichments were found for two clusters of WPG4 (WPG4-K2 which is highly expressed early, containing Man5a, CE16a and Gal28a and WPG4-K5 which was expressed late in contrast to *P. radiata* and containing OxaD and Xyl10a), however no clusters can be directly compared across treatments. Two clusters in WPG24 also have enrichment for GO terms related to hydrolase activity; WPG24-K3, which is highly induced in week 3 and contains Gal28a and CE16b and WPG24-K9, which is a large group with no specific induction pattern across harvesting points, where all the other specific genes we investigated were placed. The same pattern was demonstrated for WPG37, where all specific genes, except CE16b, were placed in a large group (WPG37-K10) with no specific induction time. WPG37-10 also had enrichment for GO terms such as protein binding, oxidation-reduction processes and catalytic activity (see Table S3 for more details). Functional enrichment was only found for one other cluster of WPG37 (WPG37-K1). This cluster was enriched for functions related to salt and water stress and was highly expressed in the second harvest point, week 6. One cluster with one or a few of the same domains as in cluster WPG37-K1 could be found for all the other treatments (*P. radiata*-K1, WPG4-K6 and WPG24-K3). For the WPG24 this was induced in the first harvest point, thus earlier than for WPG37. In *P. radiata*-K1 and WPG4-K6 there were no clear pattern of induction time for clusters with these stress domains (Fig. 3; Table S3).

**Figure 3.**
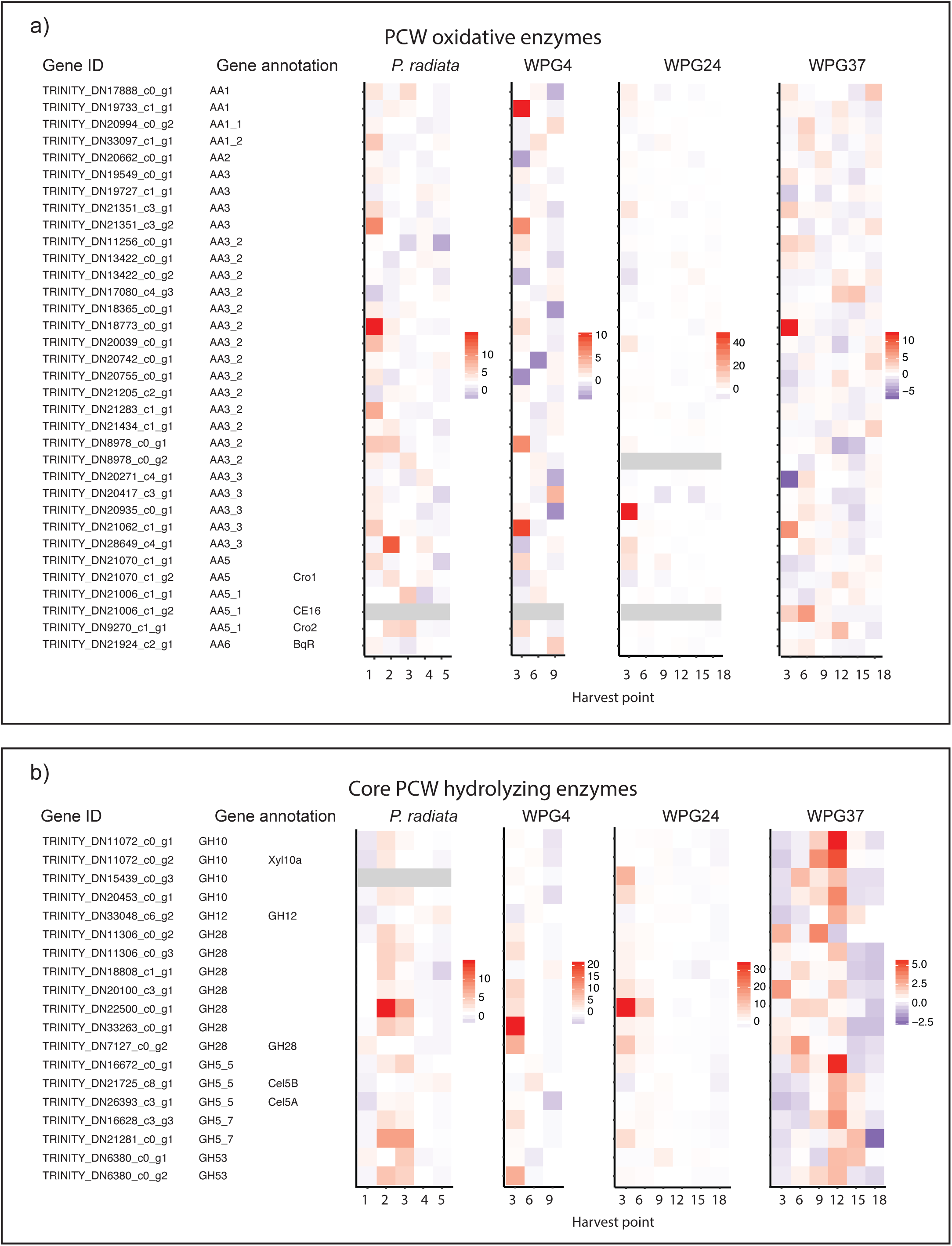
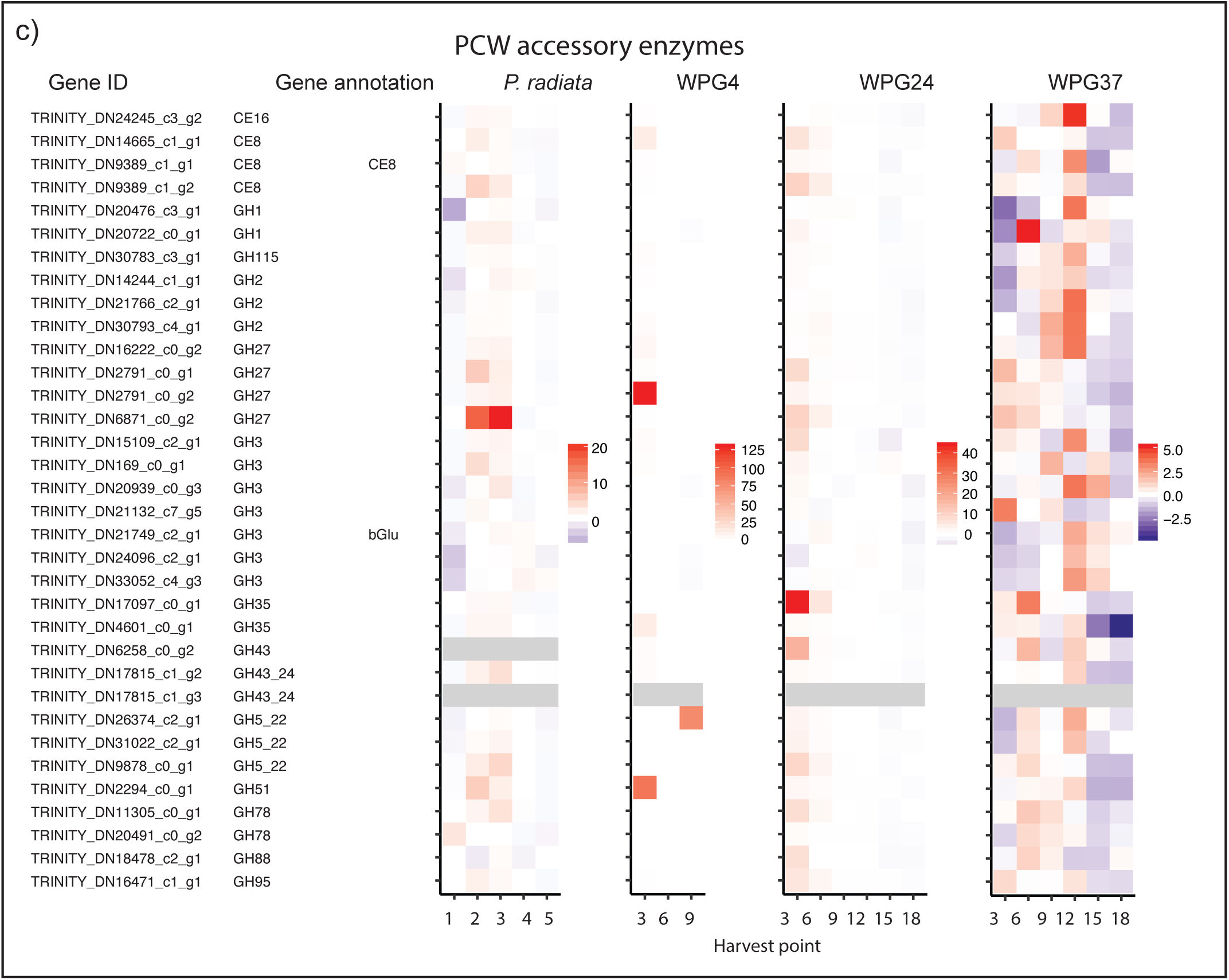
All genes were clustered into 10 groups according to their similarity in expression patterns based on read counts from RNAseq data. Each treatment was analyzed separately. The figure visualizes the relationship among these gene clusters (tree structure) and their expression pattern related to harvest point (week). Dark color indicates higher expression.

**Table 2.**
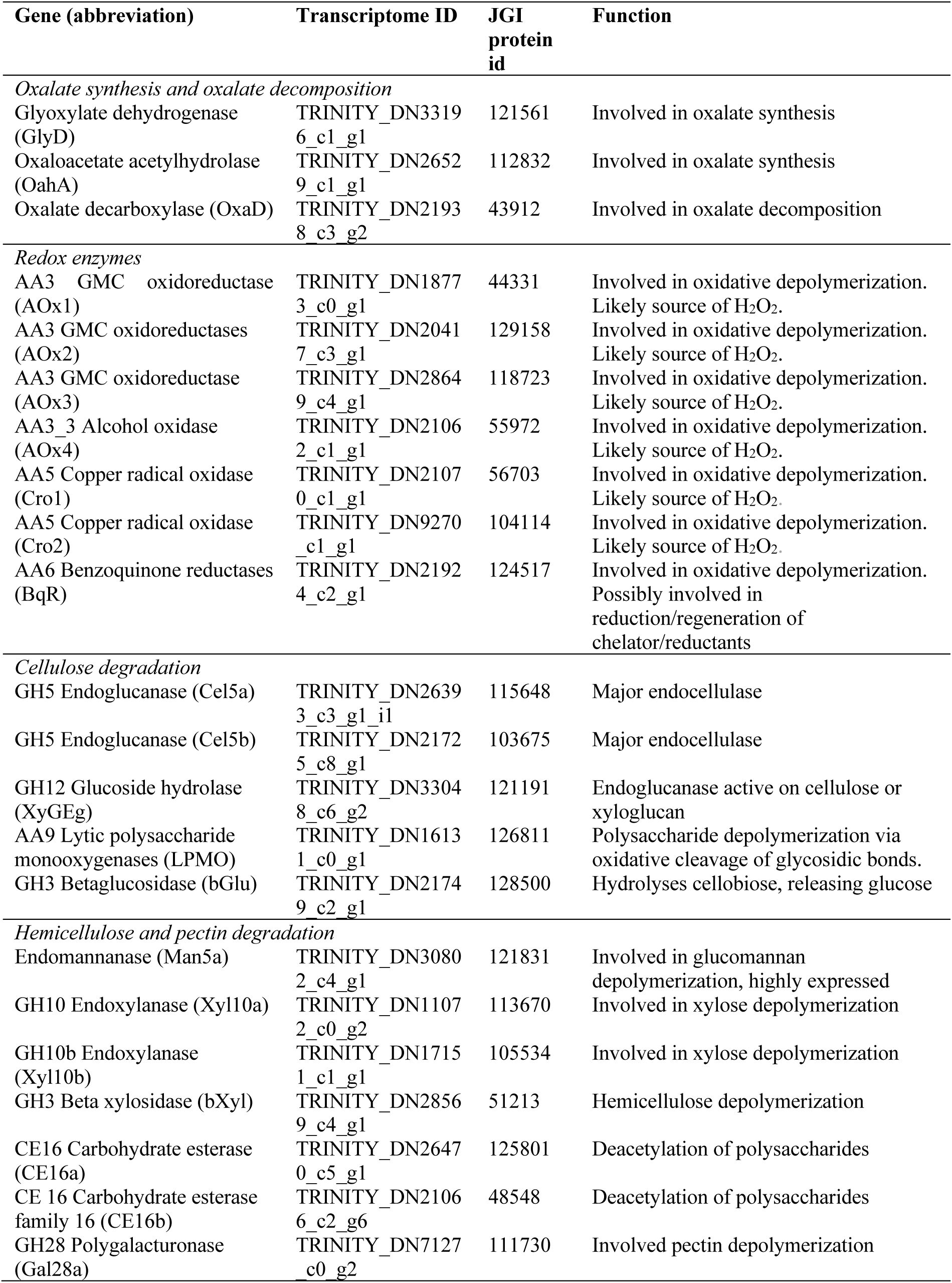
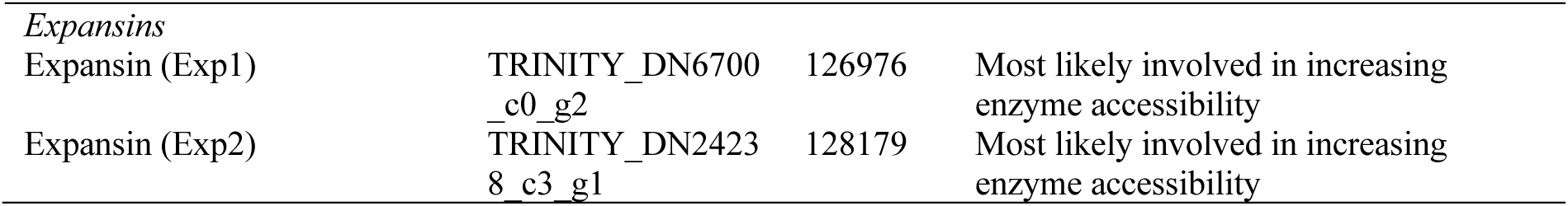
: Specific genes with functions related to plant cell wall decay investigated specifically in this study. Both for qRT-PCR analyses, raw count plots of RNAseq and their placement in cluster analyses. For identification, the transcriptome ID from our study (strain FPRL 280) and the JGI protein ID (strain MAD 698-R) is used

### Differential gene expression and functional enrichment within treatments

Pair-wise comparisons of different harvest points within treatments revealed large gene expression differences between the first and the later harvest points, for all treatments. In the *P. radiata* experiment, 2450 *R. placenta* genes were differential expressed between week 1 and week 2, while only 10 genes were differential expressed between week 2 and week 3 (Table S5).

The furfurylated wood modification demonstrated a similar temporal pattern of DE genes as the *P. radiata* experiment (Table S6). For WPG4 there were 103 DE genes between week 3 and week 6 and 481 DE genes between week 3 and week 9. No DE genes were observed between week 6 and week 9. For WPG24, large numbers of DE genes were observed between week 3 and the later time points, but again very few in between the later harvest points. For WPG37, few genes were differentially expressed (between harvest points) in general. However also here, more DE genes were found between week 3 and the later harvest points, and fewer among the later harvest points.

Functional enrichment analyses of the pair-wise analyses conformed with the expected two-step decay in this system (Table S7). These results demonstrated that for *P. radiata* experiment there was an increased expression of functions related to hydrolase and catalytic activity from week 1 to the later harvest points, especially pronounced in week 2 and week 3 (Table S7). This was also found, but to a lesser extent in WPG4 and WPG24. In WPG4, most of this response was found between week 3 and week 9, while for WPG24, the response mainly started in the comparison between week 3 and week 12 to week 18. For WPG37, this was not pronounced, and no hydrolase activity was enriched. In the first harvest points of the *P. radiata*, WPG4 and WPG24 treatments we found less enrichment of upregulated functions with the exception of some upregulation of GO terms related to protein binding and metal ion binding. However, in WPG37 there was an enrichment of several terms related to iron binding, heme binding, oxidoreductase activity and cytochrome P450s in the upregulated gene set of week 3 compared to week 18. Significant enrichment of a GO term of oxidation-reduction process was found in later stages in WPG4 and WPG24.

As week 1 in *P. radiata* was considered the first initial step of decay, with only 0.8% mass loss, we compared *P. radiata* week 1 to all other time points and treatments (Table S8). As with the timeseries analyses, a signal of downregulation of protein degradation was found in all the WPG37 harvest points compared to *P. radiata*. This was seen as enrichment of protein kinases, proteasome, F-box domain and endopeptidase activity in *P. radiata*. This pattern was also found in the early harvest points of WPG24. In contrast week 1 of the *P. radiata* showed upregulation of wood decay related functions as sugar transporters, GHs, and dehydrogenases.

### Differential expression of specific genes of interest

The gene expression of annotated CAZymes that are involved in wood decay according to Floudas et al. (34) were plotted as heatmaps across treatments (Fig. 4). On *P. radiata* the core glycoside hydrolase (GH) enzymes were expressed during intermediate harvest points (week 2 and week 3), while more of the oxidative enzymes where higher expressed early (week 1). When comparing the core enzymes between *P. radiata* and WPG37, in both treatments cellolytic activity was turned off in the late decay stages (*P. radiata* in week 4 and WPG37 in week 15). The oxidative enzymatic apparatus appears to be turned on for longer and does not have a clear induction time in WPG37 compared to *P. radiata*.

**Figure 4.**
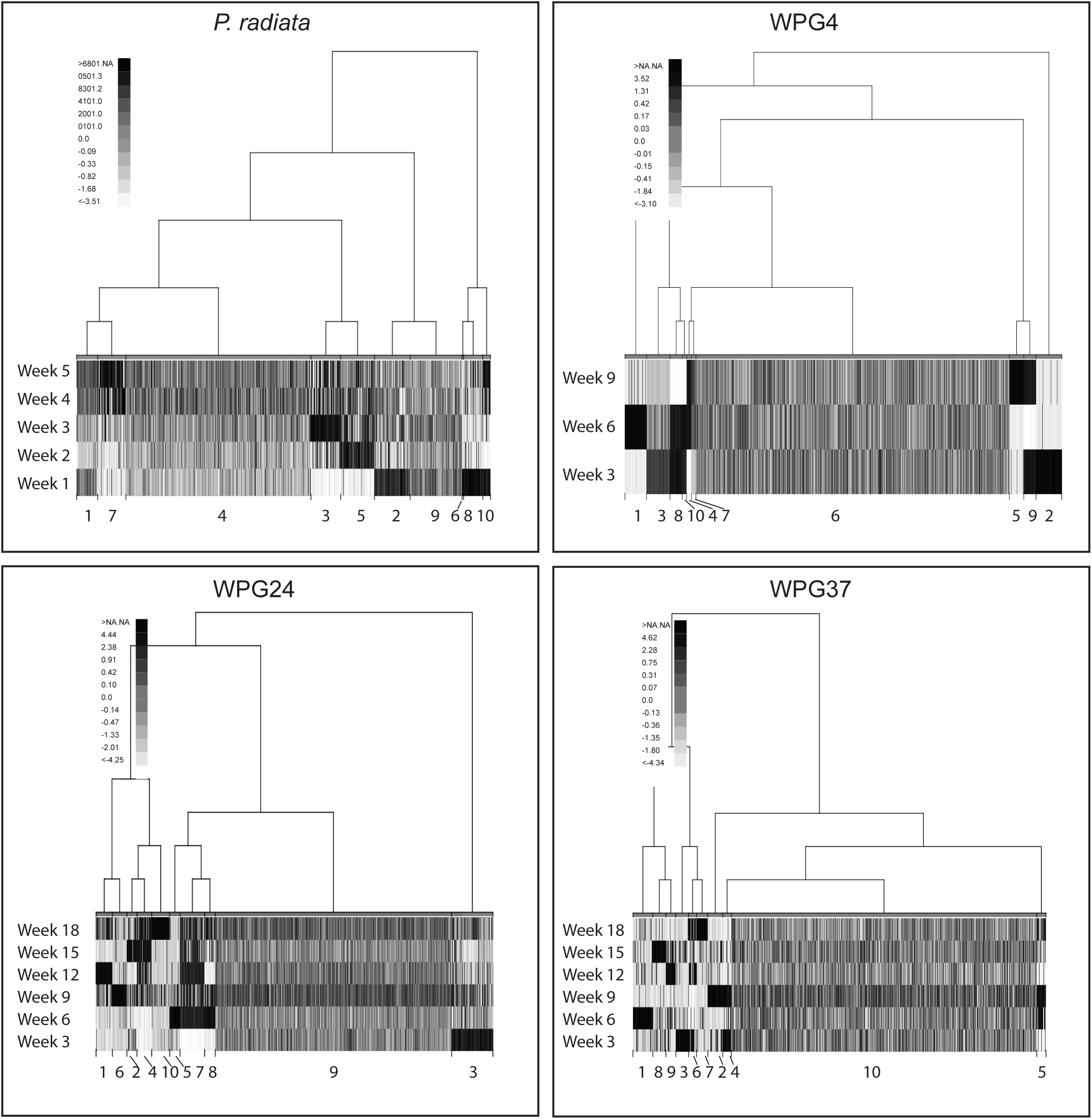
Heatmaps based on dbcan2 annotations of CAZymes suggested to be involved in plant wall decay the mean of all replicates from all experiments of *Rhodonia placenta* grown on *Pinus radiata* and different levels of modification by furfurylated *P. radiata*. Each experiment is plotted separately, with an independent scale. The Gene ID for all transcripts, the dbcan2 annotations and the corresponding gene ID from the qPCR are listed. a) the oxidizing enzymes, b) the core hydrolyzing enzymes, c) the accessory enzymes suggested to support the core enzymatic apparatus. In the accessory enzyme plot, three genes were removed from the heatmap because of extremely high expression, hiding the signal of the other genes in the *P. radiata* plot. See Fig. S2 for plot over all genes.

### Detailed qRT-PCR analyses of key genes of interest

Large RNAseq datasets tend to contain a lot of biological variation as seen in our PCA plot, which will affect downstream differential expression analyses. qRT-PCR serves both as a RNAseq control but also provides deeper/more detailed insight into the expression of individual genes. Here we have used qRT-PCR primers for key genes involved in wood decay and applied a traditional qRT-PCR approach (Thus the same genes as those reported in the cluster analyses; Table 2; Table S4; Table S9). Plots of both RNAseq and qRT-PCR data for selected genes are provided (Fig. S3-S7). The results below are based on these qRT-PCR data, RNAseq data are commented on when the trends deviated from the qRT-PCR data. It is important to keep in mind that RNAseq and qRT-PCR data are from different wood plugs (n = 4 for both datasets), i.e. the same samples are not used for both analyses. Wood samples for each experiment were selected to keep the variation small within experiments. This caused some variation between the experiments. The main effect believed to be reflected in the results is a slightly faster decay for qRT-PCR WPG37 samples than for the RNAseq WPG37 samples. It is also worth to keep in mind that qRT-PCR included an additional harvest point (week 21).

### Genes involved in oxidative depolymerization: Oxalate synthesis and oxalate decomposition

Oxalic acid is assumed to play an important role as an iron chelator and a phase transfer agent in the CMF system (16, 35). The selected genes involved in *R. placenta* oxalic acid metabolism included glyoxylate dehydrogenase and oxaloacetate dehydrogenase related to oxalic acid synthesis (GlyD, and OahA; Table 2) and oxalate decarboxylase related to oxalic acid degradation (OxaD; Table 2; Fig. S3). GlyD was upregulated in *P. radiata* compared to furfurylated samples (due to the high expression in *P. radiata* at the first harvest point). For *P. radiata* and WPG4 GlyD and OahA were upregulated the first harvest points. For WPG24 and WPG37 it seemed to be a delay in expression of OahA, showing an upregulation at the last harvesting points.

The expression levels of OxaD in qRT-PCR were low in the present study (or not detectable) and no statistically significant trends were found. The RNAseq data showed a strong induction at early-intermediate stages in the different experiments. The comparison with the lowly expressed qRT-PCR data should be interpreted with caution, but the general trends were similar (Fig. S3).

### Genes involved in oxidative depolymerization: Redox enzymes

Extracellular peroxide generation is a key component of oxidative lignocellulose degradation. The selected genes illustrated in Fig. S4 assumed to be involved in the early oxidative stage of *R. placenta* decay included three GMC oxidoreductases (CAZy family AA3) (AOx1, AOx2, AOx3 and AOx4; Table 2), two copper radical oxidases (Cro1, Cro2; Table 2) and a benzoquinone reductase (BqR; Table 2).

GMC oxidoreductases are flavoenzymes that oxidize a wide variety of alcohols and carbohydrates, with concomitant production of hydrogen peroxide or hydroquinones (36). AOx1 and AOx2 were in the present study upregulated in WPG37 compared to *P. radiata*, and for AOx2 WPG24 was also upregulated. AOx3 was highly expressed but no significant differences in expression were found within (except an upregulation WPG24 week 3) or between treatments. AOx4 showed low expression levels.

Copper radical oxidases (CAZy family AA5) are widely distributed among wood decay fungi (37), and oxidize a variety of substrates and produce H_2_O_2_. Cro1 and Cro2 were in the present study upregulated week 1 for *P. radiata* and week 21 for WPG24. Since RNAseq was not performed beyond week 18 this trend was not confirmed. Cro2 was upregulated in WPG24 compared to WPG4.

Benzoquinone reductases (CAZy family AA6) are intracellular enzymes suggested to have a role in degrading lignin and derived compounds in white rot fungi but also suggested to provide protection from oxidative stress by acting as redox active toxins and also contribute to oxidative depolymerization of wood cell wall polymers via the production of hydroquinone chelators (38). In the present study BqR was upregulated in later stages of decay both for *P. radiata* and for WPG37.

### Hydrolytic enzymes involved in polysaccharide depolymerization: Hemicellulose and pectin degradation

Fig. S5 illustrates the selected genes involved in hemicellulose hydrolysis. One endomannanase in CAZy family GH5 (Man5a; Table 2) was found to be upregulated in *P. radiata* versus WPG37. All treatments showed downregulation with increasing incubation time.

For the two endoxylanases in CAZy family GH10 (Xyl10a and Xyl10b; Table 2) there was a tendency for upregulation in furfurylated wood at 3 weeks incubation. For RNAseq an upregulation in *P. radiata* week 2.

Betaxylosidase in CAZy family GH3 (bXyl, Table 2) hydrolyses 1,4 beta D-xylans and xylobiose. bXyl was upregulated in WPG37 compared to WPG4. In *P. radiata* there was an upregulation week 3 compared to week 5, while for furfurylated wood there was an upregulation in WPG24 and WPG37 at the last harvest point (week 21).

Carbohydrate esterases catalyze the de-O or de-N-acylation of substituted saccharides. The selected carbohydrate esterase CE16a (Table 2) was upregulated in *P. radiata* compared to WPG4 and WPG37. All treatments showed downregulation with increasing incubation time. The carbohydrate esterase CE16b (Table 2) provided very low expression levels with qRT-PCR (especially *P. radiata* and WPG4). A clearer trend was found with RNAseq, especially for WPG24 and WPG37 with decreasing expression levels with increasing incubation time.

Polygalacturonase (Gal28a; Table 2) hydrolyses the alpha-1,4 glycosidic bonds between galacturonic acid residues (pectin). Gal28a was upregulated in *P. radiata* compared to WPG4 and WPG37. All treatments showed downregulation with increasing incubation time.

### Hydrolytic enzymes involved in polysaccharide depolymerization: Cellulose degradation

The endoglucanases Cel5a and Cel5b (Table 2) cause chain breaks in amorphous cellulose. Cel5a was upregulated in WPG4 vs. WPG37, and Cel5b was upregulated in *P. radiata* and WPG4 vs. WPG24 and WPG37 (Fig. S6). The expression in *P. radiata* increased with increasing incubation time.

Glucoside hydrolase XyGEg (Table 2) shows high sequence similarity to several endoglucanases with activity on cellulose and xyloglucan. This enzyme was upregulated in WPG4 vs. WPG37. In *P. radiata* samples it was a gradual increase in XyGEg with increasing incubation time.

GH3 Betaglucosidase (bGlu; Table 2) release glucose by hydrolysis of cellobiose and was upregulated in *P. radiata* and WPG4 vs. WPG37. In WPG24 and WPG37 there was an upregulation at week 9 vs. week 21.

Lytic polysaccharide monooxygenases (CAZy AA families 9, 10, 11, 13, 14 and 15) are important enzymes in lignocellulose depolymerization, that cause oxidative chain breaks in crystalline and amorphous polysaccharides. We selected a cellulose active AA9 (LPMO; Table 2). For qRT-PCR no statistically significant trends were detected for this gene except for an upregulation week 3 in WPG37.

### Expansins with a possible role in loosening plant cell-wall interactions

Expansins are hypothesized to increase enzyme accessibility by loosening plant cell-wall interactions, although no catalytic mechanism is known (39-41). The two selected genes predicted to encode expansins (Exp1 and Exp2; Table 2) are given in Fig. S7. The expression patterns of the two expansins did not show a clear trend, but for Exp1 in WPG37 the expression was upregulated week 3 and 9 compared to later decay stages.

## DISCUSSION

We have investigated if wood modification by furfuryl alcohol causes altered decomposition response in *Rhodonia (Postia) placenta*, using RNAseq and qRT-PCR, in unmodified and furfurylated *P. radiata* wood.

From our four treatments (*P. radiata*, and the three levels of modification with furfuryl alcohol polymer) we found that WPG4 closely followed the trend on *P. radiata,* while WPG24 seemed to be an accelerated version of the decay pattern of WPG37. For simplicity, we mainly focus on *P. radiata* decay processes and the comparison of *P. radiata* decay vs. WPG37 decay in the discussion.

In our *P. radiata* decay experiments we observed wood decay from incipient stages with mean mass loss of 0.8% in week 1 to severely decayed wood with a mean mass loss of 29% at week 5. Our results support the previously suggested two-step decay mechanism of brown rot fungi (13, 16, 17, 19). Week 1 of *P. radiata* demonstrated an early decay transcriptome response, with higher expression of oxidative enzymes, polygalacturanase GH28 and oxalate synthesis genes in both the RNAseq data and for the specific genes subject to qRT-PCR (Fig. 4; Fig. S4).

This early response was followed by a polysaccharide depolymerization stage, with upregulation of hemicellulases, e.g. endoxylanases (Xyl10b and Xyl10a), beta xylosidase (bXyl) and endomannanase (Man5a) in week 2 and week 3. In our study there was an abrupt downregulation of many core hemicellulases in week 4 during the later stages of decay (Fig. 4, Fig. S6); Xyl10a, Xyl10b, bXyl and Man5a). This is in agreement with observations made by Zhang and Schilling (19). It has been demonstrated that soluble sugars, in particular cellobiose, acts as the main inducing agent in the switch from oxidative (week 1) to hydrolytic depolymerization (19, 42).

Cellulose active enzymes were also induced in week 2, and were further upregulated in week 3. This was particularly obvious from the qRT-PCR expression pattern of the GH3 betaglucosidase (bGlu), and the LPMO (Fig. S6). Some cellulases are also highly expressed in the more advanced stages of decay, e.g. both the endoglucanase GH5 (Cel5b) and the glucoside hydrolase GH12 (Cel12a) are highest expressed in week 5. This high expression of cellulases at late decay stage was also observed by Zhang and Schilling (19). Thus, when the more easily accessible hemicelluloses have been degraded, expression of cellulases Cel5b and Cel12a are maintained at a high level, well into the phase where the fungus is experiencing starvation (discussed further down).

When the decay of the untreated *P. radiata* was compared to the modified wood, we observed that the oxidative enzymes were highly expressed in the first harvest point (week 3) also for the modified WPG37. This is confirming the finding in Alfredsen et al. (28) where *R. placenta* expressed high levels of oxidative enzymes and produced oxalate during 8 weeks colonization of furfurylated Scots pine. However, the pattern of repression of these oxidative processes and the oxalate synthesis found in *P. radiata* at later stages is less clear (Fig. 4). This is particularly obvious when comparing the expression of oxalate decarboxylase (OxaD) and AA3 GMC oxidoreductase (AOx1), which showed a distinct time of induction on *P. radiata* but not on WPG37 where elevated expression was randomly distributed over time between samples. This can conceivably be explained by a repeated exposure to regions with heavily modified substrate, thus re-inducing oxidative processes to overcome furfurylation in order to expose degradable substrate to the fungus allowing further growth, or a direct response to the furfuryl polymer.

However, the strong induction of core hydrolyzing enzymes and accessory enzymes observed in week 2 and 3 in *P. radiata* seemed comparable to WPG37 from week 6-12, and partially week 15 as observed by the RNAseq CAZymes analyses and the qRT-PCR analyses (Fig. 4, Fig. S6). Thus, the switch to turn on the core hydrolyzing enzymes in the intermediate decay stages was observed in both untreated and furfurylated wood in our experiments, thus we hypothesize that furfurlyation does not influence the availability of these soluble sugars to such an extent that it inhibits induction. Previous studies have showed that *R. placenta* grown on wood modified with both furfurylation and acetylation show similar or decreased levels of core wood hydrolyzing enzymes (26, 28). Our study, with a longer time series, does not suggest reduced expression of these genes, but rather a delayed and elongated process. The elongated process could also be the reason for the placement of all the specific plant cell wall related genes in the large cluster with no obvious induction time in WPG37. The longer incubation times used in this study, and the elongated processes can explain why this trend was never observed in previous studies. Thus, based on the observation that a delayed but, similar pattern was observed in both the *P. radiata* and WPG37 experiments, we conclude that the furfurylation does not directly influence the expression of these core PCW degrading enzymes.

There are several hypotheses regarding the mode of action of modified wood against brown rot decay fungi (43, 44), summarized in (45). Briefly they hypothesize that: unavailability of easily accessible nutrients (43, 44), enzyme non-recognition (7), micropore blocking (46), and moisture exclusion due to OH-group blocking/reduction (44), and/or reduction in void volume may be important modes of action (47, 48). In our study we have proven that the enzymes do recognize the substrate in the modified treatments, thus we can rule out enzyme non-recognition. The main conclusion from Ringman et al. (45) was that “only one theory provides a consistent explanation for the initial inhibition of brown rot degradation in modified wood, that is, moisture exclusion via the reduction of cell wall voids. Other proposed mechanisms, such as enzyme non-recognition, micropore blocking, and reducing the number of free hydroxyl groups, may reduce the degradation rate when cell wall water uptake is no longer impeded”. The delay in gene expression in our study can be explained by initial blocking of wood polysaccharides by furfuryl alcohol polymers compared to unmodified wood. In addition, a further delay in decay of modified wood versus *P. radiata* is expected due to lower wood moisture content in the modified substrate. Moreover, the strong enrichment of salt stress genetic functions in the WPG37-K1 clustering that was not found in the other treatments may support the conclusions by Ringman et al. (45) that moisture exclusion may be an important mode of action in this modification method.

As a support for the notion that the furfurylation is non-toxic for the fungus, we found no evidence for expression of more defense mechanisms in the modified wood. *Rhodonia placenta* is known to be highly tolerant to substances as copper, mainly due to the ability to produce oxalic acid that chelate and precipitate copper and other transition metals (49). However, in addition to these more general functions that are difficult to separate from wood decay mechanisms, copper tolerant fungi are also known to express catalases and ATP-pumps related to copper transport in response to toxins (50). These functional categories were not enriched in the differential expressed gene sets in our study when comparing WPG37 to the *P. radiata*, indicating that the fungus is not experiencing a more toxic atmosphere in the modified wood.

Notably, the most pronounced transcriptome differences in the *P. radiata* compared to the modified wood is the strong induction of functions related to the ubiquitin/proteasome pathway, protein degradation and the RAS pathway. All these functions are related to carbon starvation and are highly enriched in the late decay of the unmodified *P. radiata*. Thus, we hypothesize that *R. placenta* growing on *P. radiata* is starving and that the fungus has consumed the majority of available carbon sources in the wood in the latest harvest points of the *P. radiata* experiment. This is further supported by the downregulation of CAZymes the last two weeks in this experiment.

The ubiquitin/proteasome pathway is a conserved pathway in all eukaryotic organisms, and is important to various cellular processes as recycling of intracellular protein (where unnecessary proteins are degraded to amino acids that can be reused to produce new proteins) and programmed cell death. Recently, the pathway was shown to be activated by carbon starvation in the ectomycorrhizal basidiomycete species *Paxillus involutus* (51). In *P. involutus*, 45% of the transcripts were differentially regulated during carbon starvation. This large response is also shown in our study where most enriched functions in *P. radiata* compared to the WPG37 for the time series analyses could be connected to the U/P or the Ras pathway. This could reflect the higher need for translocation of nitrogen rich resources in the *P. radiata* substrate to areas that need new protein synthesis than is the case in WPG37 or simply reflect a general starving response due to lack of substrate.

The other pattern observed as upregulated in *P. radiata* compared to WPG37 was several domains related to Ras proteins. These proteins are involved in cell proliferation and growth of the mycelia. Carbon starvation supports an upregulation of these proteins. In starving mycelia it has been shown that the diameter of the hypha is reduced, while it grows to cover larger area (51, 52). Ras proteins have been shown to enhance the formation of pseudohyphal growth in starving yeast cultures. These pseudohyphae are thin and long cells extending away from the culture, searching for nutrients (53). Ras proteins might well be involved in a change in growth to search for more nutrients in starving mycelia across the fungal kingdom.

In all our experiments we have investigated one strain, the FPRL 280. Thaler *et al.* (54) suggested significant differences in the regulation of key lignocellulose degrading enzymes between the previously sequenced MAD-698-R strain and the FPRL 280 strain used in this work. This proposed difference is supported by the current study with a mapping success of only 40-60% when the reads of European strain FPRL 280 were mapped to the genome of MAD 698-R. The difference might have been present since time of isolation or (less likely) the changes might have occurred during storage of these strains. This highlights the importance of providing verifiable strain information when publishing decay studies and when comparing American versus European wood decay testing.

Furthermore, for decay testing, the sample size, sample geometry and wood anatomy influence the colonization rate. Sampling time and extent of decay is crucial when examining gene expression or secretome of fungal colonization of wood. The studies by Zhang et al. (17) and Zhang & Shilling (19) used thin wafers and harvested at different distances from the *R*. *placenta* hyphal front. This wafer approach works relatively well to separate different initial decay stages. For more advanced stages of decay a different approach is needed. In the current study we selected small and homogeneous (earlywood) samples in order to enable fast colonization and as homogenous solid wood substrate for decay as possible. Still, some variation of gene expression is expected as new colonization pathways will be found close to areas with previously colonized wood. This effect is expected to increase with increasing WPG level, since areas long colonized will cease to provide nutrients able sustain survival while the others more recently invaded will provide nutritious substrates. In our experiments, the wood modification treatment modifies the composition of the wood, and the fungus is therefore forced to respond differently than when it encounters unmodified wood. Hence, comparison at similar mass losses between treatments was not the goal of this study, but rather the shift of gene expression over time between and within the different treatments.

Increased knowledge about brown rot decay mechanisms is important for an expanded understanding of the fungal decay process in general since ecosystem carbon flows are closely linked to wood degrading fungi. From an industry perspective the findings in this study show that successful inhibition of the initial oxidative decay is a clue to success for future wood protection systems.

## CONCLUSIONS

This is the first time the entire gene expression pattern of a decay fungus is followed in untreated and modified wood from initial to advanced stages of decay. From these observations, we have demonstrated how the furfurlyated modification delays the fungus gene expression while growing on this substrate. All treatments (modified and unmodified) expressed the expected staggered decay mechanism, i.e. oxidative enzymes earlier, and then the hydrolyzing enzymes later. Thus, we show that the fungus experiences a similar substrate situation on the modified wood, or at least is following the same modality of gene expression. The major difference is that the responses are delayed and elongated compared to unmodified wood. However, from the downregulation of CAZymes in the latest harvest points in the WPG37, the fungus seems to be finalizing the decay also in the furfurylated wood at a mass loss of only 14%, compared to 29% in *P. radiata*. This suggests that the fungus never get access to the remaining carbohydrates from the modified wood even at low mass loss. The lower levels of modification in the WPG4 and WPG24 treatments show intermediate expression differences and higher mass loss. The mode of action of the modification is still uncertain. However, the similar decay mechanisms, and the lack of expression of defense mechanisms may indicate that the furfurylation functions as both a physical barrier and a factor that creates a less hydrated environment for the fungus. These hypotheses should be investigated to further improve environmentally friendly modification processes.

## MATERIAL AND METHODS

### Wood material

In order to get as homogeneous wood material as possible plugs (Ø = 6 mm, h = 10 mm) were prepared from *Pinus radiata* D. Don earlywood according to Beck et al. 2017 (55). The boards were provided by Kebony ASA, Skien, Norway. Before treatment all samples were dried at 103 °C for 18 hours then cooled down in a desiccator before initial dry weight was recorded. The furfurylation process was performed with three different concentrations of furfuryl alcohol synthesis grade > 98% (Merck, Darmstadt, Germany) according to the formula by Kebony with; furfurylalcohol to water ratio of 7:10 (WPG37) the commercial treatment level, 4:10 (WPG24) and 1:10 (WPG4). The samples were left soaked in the furfuryl alcohol solutions for 15 days. Sets of five samples were wrapped in aluminum foil and cured at 120 °C for 16.5 hours. All samples including *P. radiata* were leached according to EN 84 (1997) as a pre-weathering test and dried at room temperature. In order to provide Weight Present Gain (WPG) and initial dry mass after treatment the samples were dried at 103 °C for 18 hours then cooled down in a desiccator before the dry weight was recorded. The samples were left in a climate chamber at 65% relative humidity and 20 °C until stable weight before they were wrapped in sealed plastic bags and sterilized by gamma irradiation (25 kGY) at the Norwegian Institute for Energy Technology.

### Decay test

The brown rot fungus in this experiment was *Rhodonia placenta* (Fr.) Niemelä, K.H. Larss. & Schigel (syn. *Postia placenta*) strain FPRL 280. The fungus was first grown on 4% (w/v) Difco™ malt agar (VWR) media and plugs from actively growing mycelia were transferred to a liquid culture containing 4% (w/v) Difco™ malt (VWR). After two weeks the liquid culture was homogenized with a tissue homogenizer (Ultra-turrax T25, IKA Werke GmbH & Co. KG, Staufen, Germany).

A modified E10-16 soil-block test (21) was used. Agar plates (TC Dish 100, standard, Sarstedt AG & Co., Nümbrecht, Germany) (Ø = 87 mm, h = 20 mm) containing soil (2/3 ecological compost soil and 1/3 sandy soil) was adjusted to 95% of its water holding capacity according to ENV 807 (CEN 2001). A plastic mesh was used to avoid direct contact between the samples and the soil. A 300 µl inoculum of homogenized liquid culture was added to each sample. Eight samples of the same treatment were added to each plate and four replicate plates were used.

Samples were incubated at 22°C and 70% RH and harvested every third week. Fungal mycelium was manually removed from the wood surface with Delicate Task Wipes (Kintech Science, UK). Eight samples from each treatment and each harvesting point were dried at 103°C for 18 hours in order to provide data for mass loss (mean WPG: WPG4 3.8±0.7, WPG24 24.6±4.1, WPG37 36.1±5.5). The remaining samples were wrapped individually in aluminum foil and put directly into a container with liquid nitrogen. The samples were then stored at -80°C.

For RNAseq analyses samples from the following harvesting points were used: *P. radiata* week 1, 2, 3, 4 and 5, WPG4 week 3, 6 and 9, WPG24 week 3, 6, 9, 12, 15 and 18. For qRT-PCR samples from the following harvesting points were used: *P. radiata* week 1, 3 and 5, WPG4 week 3, 6 and 9, WPG24 week 3, 9, 15 and 21, WPG37 week 3, 9, 15 and 21.

### RNA purification and cDNA synthesis

Wood powder from frozen samples was obtained by cutting the plugs into smaller pieces with a garden shears wiped with 70% alcohol and thereafter Molecular BioProducts™ RNase AWAY™ Surface Decontaminant (Thermo Scientific, Singapore). The wood samples were immediately cooled down again in Eppendorf Tubes™ in liquid nitrogen. Fine wood powder was produced in a Retsch 300 mill (Retsch mbH, Haan, Germany). The wood samples, the 100-mg stainless steel beads (QIAGEN, Hilden, Germany) and the containers were chilled with liquid nitrogen before grinding at maximum speed for 1.5 min. They were then cooled in liquid nitrogen again before a second round of grinding.

For Illumina seq MasterPure™ Plant RNA Purification KIT (epicentre, Madison, WI, USA) was used according to the manufacturer’s instruction for total RNA extraction with 100 mg wood sample. Mean WPG for the four replicate samples: WPG4 3.6±0.5, WPG24 24.8±1.2, WPG37 39.7±2.2.

For qRT-PCR MasterPureTM Complete DNA and RNA Purification KIT (epicenter, Madison, WI, USA) was used according to the manufacturer’s instruction with 90 mg wood sample. (A different kit was used because the kit used for Illumina analyses was no longer produced). Mean WPG for the four samples: WPG4 4.1±0,4, WPG24 20.9±0.9, WPG37 33.7±3.8. NanoDrop™ 2000 spectrophotometer (Thermo Scientific, Singapore) was used to quantify RNA in each sample. To convert RNA to cDNA TaqMan Reverse Transcription Reagent KIT (Thermo Scientific, Singapore) was used according to the manufacturer’s instructions. Total reaction volume was 50 µl. 300 ng RNA were reacted with oligo d(T)_16_ primer in RNase free water (Qiagene, Hilden, Germany). The solution was incubated two cycles in the PCR machine (GeneAmp^®^ PCR System 9700, Applied Biosystems, Foster City, CA, USA) at 65 °C/5 min and 4 °C/2 min. The PCR machine was paused and the master mix added. The next three cycles included 37 °C/30 min, 95 °C/5 min and 4 °C/indefinite time. In addition to the test samples, two samples without RNA were added as controls and used for each primer pair. After the cDNA synthesis, 50 µl RNase free water (Qiagene, Hilden, Germany) was added to the samples and mixed well.

### RNAseq sequencing, quality control and trimming

All Illumina libraries and sequencing were performed by the Norwegian Sequencing Centre (http://www.sequencing.uio.no). All samples were prepared with the strand specific TruSeq™ RNA-seq library preparation method. In order to produce a high quality *de novo* transcriptome we sequenced the strain grown on a 4% (w/v) Difco™ malt agar (VWR) media in addition to ten selected samples spanning the entire experimental setup on one lane on the Hiseq2500 producing 125 bp paired end sequences. These libraries generated ~395 million read pairs, which after quality control and trimming was reduced to ~250 million read pairs.

For the experiment itself, we collected 80 samples of which three failed library preparation resulting in 77 NextSeq samples in total. The sequenced libraries generated ~28 million trimmed reads on average.

The quality was evaluated using FastQC v. 0.11.2 (56) and trimming performed with Trimmomatic v. 0.36 (57) with the following parameters: TruSeq3-SE.fa:2:30:10 MAXINFO:30:0.4 MINLEN:30 for the NextSeq samples. For the Hiseq2500 samples, the built-in trimmomatic option in Trinity v. 2.2.0 (58) with the following parameters: TruSeq3-PE.fa:2:30:10 SLIDINGWINDOW:4:5 LEADING:5 TRAILING:5 MINLEN:30 MAXINFO:30:0.4 was used.

### Transcriptome assembly and evaluation

We initially attempted to use the *R. placenta* genome generated from an American *R. placenta* strain in our analysis (Postia_placenta_mad_698, full fasta sequence (soft masked) from the Ensembl fungi FTP server). The genome-based attempt using Bowtie2, Tophat2 and Cufflinks resulted in very poor mapback (~50-70 %) despite doing parameter sweeps (data not shown). We also attempted to use the genome-guided option of the *de novo* transcriptome assembler Trinity that yielded a more fragmented transcriptome compared to the clean *de novo* version (data not shown).

The final transcriptome was generated using a *de novo* strategy with the HiSeq trimmed reads and Trinity v. 2.2.0 (seqType fq, max_memory 250G, SS_lib_type RF, CPU 20, bflyHeapSpaceMax 10G and bflyCPU 20) on our local computing cluster. The Trinity bfly process failed for 110 elements which were rerun with more memory. The finished transcriptome contained 56 520 contigs (Table S1).

We also evaluated the assembly using BUSCO v. 2.0 using the recommended parameters and the corresponding fungi database created 26^th^ of January 2017 with 85 species and 290 BUSCO genes (59). Of the 290 BUSCO genes 288 were found complete (99.3 %) (Table S2).

### Assembly annotation using Trinotate and the JGI Fungi Portal

The assembly was subjected to the Trinotate annotation pipeline according to the manual which provided a generic annotation of assembly. However, as there is a study (17) looking into key wood-degrading genes using the annotation from the *R. placenta* genome, we manually annotated our assembly using the predicted proteins reported at the JGI Fungi Portal (https://genome.jgi.doe.gov/programs/fungi/index.jsf) using TBLASTN from BLAST+ (60) with an e-value cutoff 1e-50. Finally, we ended up with a custom annotation being a hybrid of the generic trinotate annotation where identifiers from the *R. placenta* genome have been added. Finally, we manually evaluated the hits from the key genes reported in Zhang *et al.* (17) to overwrite any generic annotation provided by trinotate as these have been manually verified and submitted to the CAZy database (http://www.cazy.org/).

### Mapping and abundance estimation of NextSeq samples

Mapping was performed using the built-in mapping option in Trinity v. 2.3.2 with the RSEM count estimation method, the bowtie alignment method and specifying SS_lib_type R. The counts were collected per ‘gene’ using the abundance_estimates_to_matrix.pl script in Trinity and the resulting count matrix used for differential gene expression in R.

### Differential gene expression analysis

In R v3.4.1 the overall data was initially explored using VarianceParition v1.8 (61). Differential gene expression analyses were performed using edgeR v 3.20.1 (62, 63). In addition some of the plot functions in DESeq2 v1.18.0 (64) were used to explore that overall data and to plot raw counts.

The overall workflow with code is available in the supplementary information, but briefly we followed the steps described below:

The coldata object describing the overall experiment was set up with time, treatment, mass loss and condition where condition in practice is the intercept between time and treatment. We initially filtered on counts per million (CPM, cutoff = 1 in at least 2 libraries) and set a full model consisting of treatment, time, condition and mass loss to explore the data in variancePartition. We also made heatmaps using pheatmap v1.0.8 and log2 transformed count data.

Multifactorial differential expression analysis was performed both in EdgeR and DESeq2 (the latter to enable use of various plot functions). In DESeq we opted for a ~treatment+time formula. Running a more complex design with interaction (:) was not possible as DESeq2 cannot handle partial models (time points for the different treatments do not overlap completely). Furthermore, we opted for the likelihood ratio test (LRT) which examines two models for the counts - the full model and a reduced model. This then determines if the increased likelihood of the data using the extra term(s) in the full model is more than expected.

In edgeR we opted for a ~0 + treatment+time formula without generating an intercept term. When estimating overall dispersion, we used the robust=TRUE option to better handle outliers in the data. For the multifactorial test itself we chose to replace the standard glmFit with glmQLFit which uses a quasi-likelihood F-test on the likelihood ratio statistics instead of the chisquare approximation. In this way we should obtain a more conservative control of the type I error rate as it takes into account the uncertainty in estimating dispersion for each gene - especially when the number of replicates is small.

To enable exploration of specific contrasts between given conditions we also ran a ~0 + condition formula in edgeR using the same setup as described above.

We evaluated each treatment vs the *P. radiata* over time from the treatment+time multifactorial analysis. For the pairwise contrasts between harvest time within treatments we used the ~condition multifactorial analysis.

The overall exploration of the data revealed that the WPG24 treatment series has a few outliers. RNAseq data is highly variable and the individual variation between replicates can be large. We found that 2 replicates in the WPG24 3 weeks condition were relatively deviating from the other two replicates. We reran the above described analysis without the most extreme outlier (WPG24 3 weeks replicate 2) and observed a large change in the number of reported significant differentially expressed genes. However, as these differences not necessary improved the results we decided to keep all samples in the final dataset.

### Clustering using MCDluster.seq

As an alternative to the pair-wise differential expression analyses the read count data were clustered based on similarity in expression patterns using the MBCluster.seq package in R. The EM clustering algorithm was used.

### Functional summary

Functional enrichment analysis was used to characterize function of the differential expressed genes of the various treatments and clusters. A Python script was used to perform functional enrichment analysis of PFAM domains and GO terms using Fisher’s exact test (http://cgrlucb.wikispaces.com/Functional+Enrichment+Analysis).

### CAZyme annotations

The transcripts were translated using TransDecoder (https://github.com/TransDecoder/TransDecoder/wiki). All protein sequences were annotated in dbCAN2 (65). The standardized (median/median absolute deviation) gene expression patterns of the resulting annotated transcripts were plotted using in a heatmap using R (66).

### qRT-PCR

The qRT-PCR specific primers used to determine the transcript levels of selected genes were designed with a target T_m_ of 60°C and to yield a 150 base pair product. qRT-PCR was performed using ViiA 7 by Life technologies (Applied Biosystems, Foster City, CA, USA). The master mix included for each sample: 5 µl Fast SYBR^®^Green Master Mix (Thermo Scientific, Singapore), 0.006 µl 10 µM forward primer, 0.006 µl 10 µM reverse primer, 2.88 µl RNase free water (Qiagene, Hilden, Germany) and 2 µl test sample (total volume 10 µl). The qRT-PCR run included the following stages: Hold stage with initial ramp rate 2.63 °C/s, then 95.0 °C for 20 seconds. PCR stage with 40 cycles of initial ramp rate 2.63 °C/s, 95.0 °C, ramp rate of 2.42 °C followed by 60.0 °C for 20 seconds. The melt curve stage had an initial ramp rate of 2.63 °C/s then 95.0 °C for 15 seconds, ramp rate of 2.42 °C/s 60.0 ° for one second, then 0.05 °C/s.

Two constitutive housekeeping genes, β-tubulin - βt (113871) and α-tubulin α-t (123093) were used as a baseline for gene expression. The target genes (Tg) and the endogenous controls in this study are listed in Table 1. Protein ID according to *Postia placenta* MAD 698-R v1.0 genome, The Joint Genome Institute (https://genome.jgi.doe.gov/pages/search-for-genes.jsf?organism=Pospl1). Threshold cycle values (Ct) obtained here were used to quantify gene expression.

Software used to export the Ct values was QuantStudioTM Real-Time PCR System (Applied Biosystems by Thermo Fiches Scientific, Foster City, CA, USA). Ct-value of βt, αt and Tg were used to quantify gene expression according to the following equation; 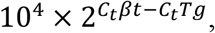 giving an arbitrary baseline expression of β-tubulin and α-tubulin of 104. As an internal control, the expression of βt and αt was compared using the same equation, showing a stable expression, with αt being expressed at approximately 80% relative to βt. Only data for βt was included in this paper.

### Statistical analysis of qRT-PCR

All statistics were performed in JMP (Version Pro 13, SAS Institute Inc., Cary, NC, USA). Significance of differences in expression levels of each gene were calculated with Tukey honest significant difference (HSD) test. A probability of ≤ 0.05 was the statistical type-I error level.

## DATA AVAILABILITY

RNA-seq data generated for this study were deposited at the Sequence Read Archive under accession XXXXXX. The transcriptome assembly and all lists of differentially expressed genes were deposited in Dryad under accession doi:XXXXXX.

## ACKNOWLEDGEMENT

Thanks to Sigrun Kolstad and Inger Heldal (NIBIO) for help with RNA purification and qRT-PCR. Thanks to Stig Lande (Kebony) for helpful discussions regarding treatment of the samples. This project was financed by The Research Council of Norway 243663/E50 BioMim.

